# Testing the neutral hypothesis of phenotypic evolution

**DOI:** 10.1101/089987

**Authors:** Wei-Chin Ho, Yoshikazu Ohya, Jianzhi Zhang

## Abstract

It is generally accepted that a large fraction of genomic sequence variations within and between species are neutral or nearly so^1^. Whether the same is true for phenotypic variations is a central question in biology^2-7^. On the one hand, numerous phenotypic adaptations have been documented^2,8,9^ and even Kimura, the champion of the neutral theory of molecular evolution, believed in widespread adaptive phenotypic evolution^1^. On the other hand, phenotypic studies are strongly biased toward traits that are likely to be adaptive^9^, contrasting genomic studies that are typically unbiased. It is thus desirable to test the neutral hypothesis of phenotypic evolution using traits irrespective of their potential involvement in adaptation. Here we present such a test for 210 morphological traits measured in multiple strains of the yeast *Saccharomyces cerevisiae* and two related species. Our test is based on the premise that, under neutrality, the rate of phenotypic evolution declines as the trait becomes more important to fitness, analogous to the neutral paradigm that functional genes evolve more slowly than functionless pseudogenes^10^. Neutrality is rejected in favor of adaptation if important traits evolve faster than less important ones, parallel to the demonstration of molecular adaptation when a functional gene evolves faster than pseudogenes. After controlling for the mutational size, we find faster evolution of more important morphological traits within and between species. By contrast, an analysis of 3466 yeast gene expression traits fails to reject neutrality. Thus, yeast morphological evolution is largely adaptive, but the same may not apply to other classes of phenotypes.

---

Analogous to the neutral hypothesis of molecular evolution^1^, the neutral hypothesis of phenotypic evolution allows the presence of purifying selection; neutrality is rejected only when positive selection is invoked. Under the neutral hypothesis, compared with traits that are relatively unimportant to fitness, relatively important traits should be subject to stronger purifying selection and evolve more slowly given the same speed of mutational input (Fig. 1a). However, if relatively important traits evolve faster than relatively unimportant traits, the neutral hypothesis would no longer hold and the only reasonable explanation would be stronger positive selection acting on relatively important traits than relatively unimportant ones (Fig. 1b). This test of phenotypic neutrality differs from previous tests^3,4,11-15^, which consider only one trait at a time and effectively require the intensity of positive selection to surpass that of purifying selection to reject neutrality. Because this requirement is sufficient but not necessary for demonstrating positive selection, it is replaced with the criterion of a positive correlation between trait importance and evolutionary rate to improve the power of the test (see Methods and Discussion).

**Figure 1.**
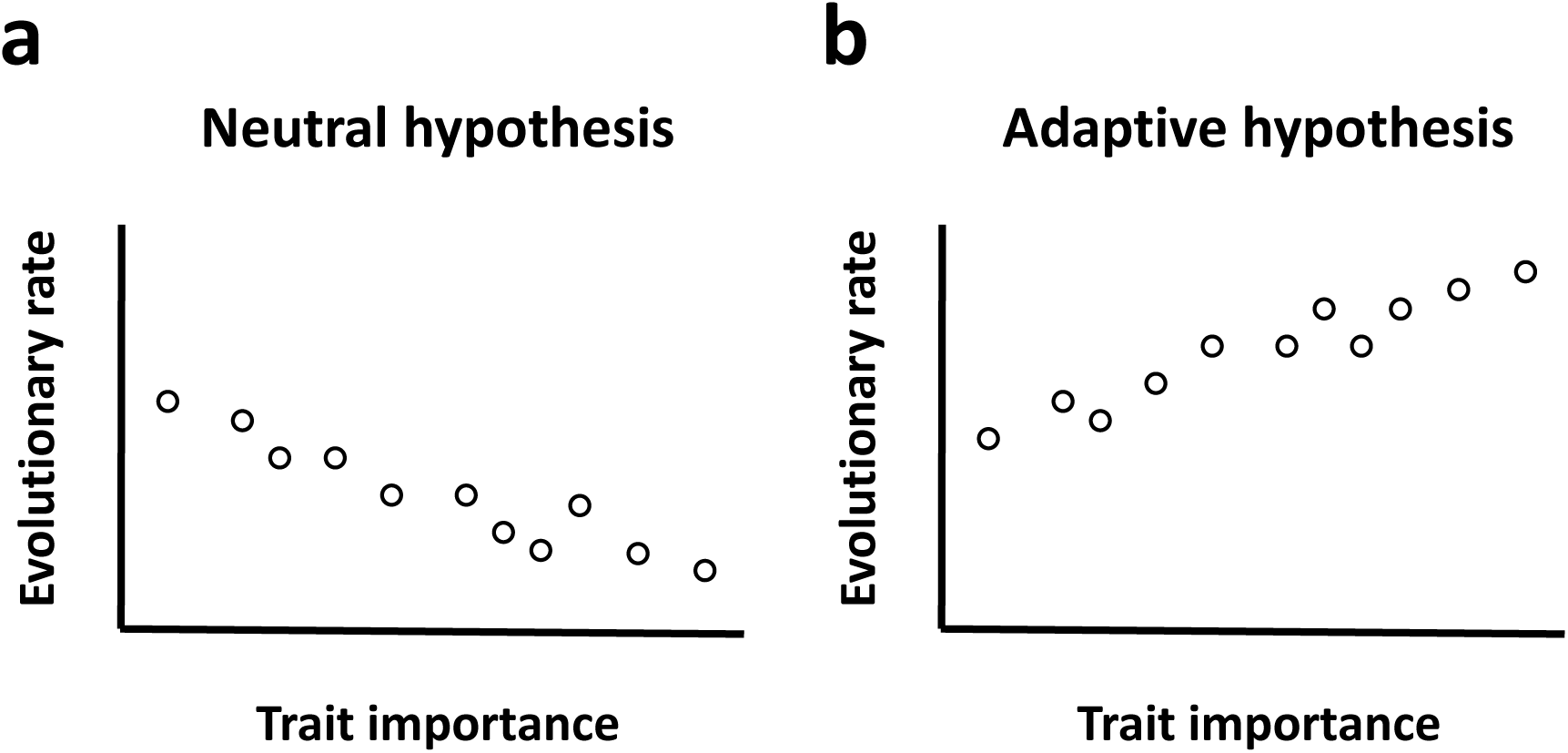
Comparing evolutionary rates among traits to test the neutral hypothesis of phenotypic evolution. **a**, Under the neutral hypothesis, relatively important traits evolve more slowly than relative unimportant traits. **b,** Higher evolutionary rate for more important traits rejects the neutral hypothesis and supports the adaptive hypothesis. Each circle represents a trait.

We first tested the neutral hypothesis of phenotypic evolution in a set of 210 morphological traits chosen purely based on the feasibility of measurement^16^ rather than potential roles in adaptation (**Supplementary Data 1**). These traits were quantified by analyzing fluorescent microscopic images of triple-stained cells^16^ from 37 natural strains of *S. cerevisiae*^17^. For a given trait, we defined the phenotypic difference between two strains as the absolute difference in their trait value, relative to their average trait value. Phenotypic differences were corrected for potential environmental heterogeneity in the measurement and sampling error to allow among-trait comparison (**Supplementary Data 2**; see Methods). We then estimated, for each trait, the mean evolutionary distance (*ED*) among all 666 pairs of the 37 strains by averaging their corrected pairwise phenotypic differences.

To test the neutral hypothesis, we used measures of trait importance (*TI*) for the 210 traits (**Supplementary Data 1**), where *TI* is 100 times the fitness effect caused by 1% change in trait value, estimated using the fitness and phenotype data of thousands of single gene deletion strains of *S. cerevisiae*^18^. We found that mean *ED* decreases with *TI* (Fig. 2a), meaning that relatively important traits evolve more slowly than relatively unimportant ones. However, the rate of phenotypic evolution is determined by both the rate of mutational input and the direction and magnitude of natural selection. The rate of mutational input for a trait is the average effect size of a random mutation on the trait (i.e., mutational size or *MS*) multiplied by the mutation rate per genome per generation. Because the mutation rate is the same for all traits, we need only consider *MS*. We estimated the *MS* for a trait by the mean phenotypic effect of 4718 individual gene deletions on the trait^18^ (**Supplementary Data 1**). Although spontaneous mutations are expected to have smaller phenotypic effects than gene deletions, it is reasonable to assume that the average phenotypic effect of spontaneous mutations on a trait is approximately proportional to that of gene deletions on the trait. In other words, *MS* can be used as a proxy of spontaneous mutational size when different traits are compared. As was previously discovered^18^, *MS* decreases precipitously with *TI* (Fig. 2a). This observation indicates that relatively important traits are affected by mutations to a smaller degree than are relatively unimportant traits, which has likely resulted from stronger selection for mutational robustness of more important traits^18^. To control the impact of mutational input on the rate of phenotypic evolution, we measured the evolutionary rate of a trait in the unit of its mutational size by dividing mean *ED* by *MS* for each trait. We found mean *ED*/*MS* to increase significantly with *TI* (Fig. 2b), suggesting that relatively important traits evolve faster than relatively unimportant traits in the unit of mutational size. This finding is inconsistent with the neutral hypothesis of phenotypic evolution (Fig. 1a), but supports the adaptive hypothesis (Fig. 1b).

**Figure 2.**
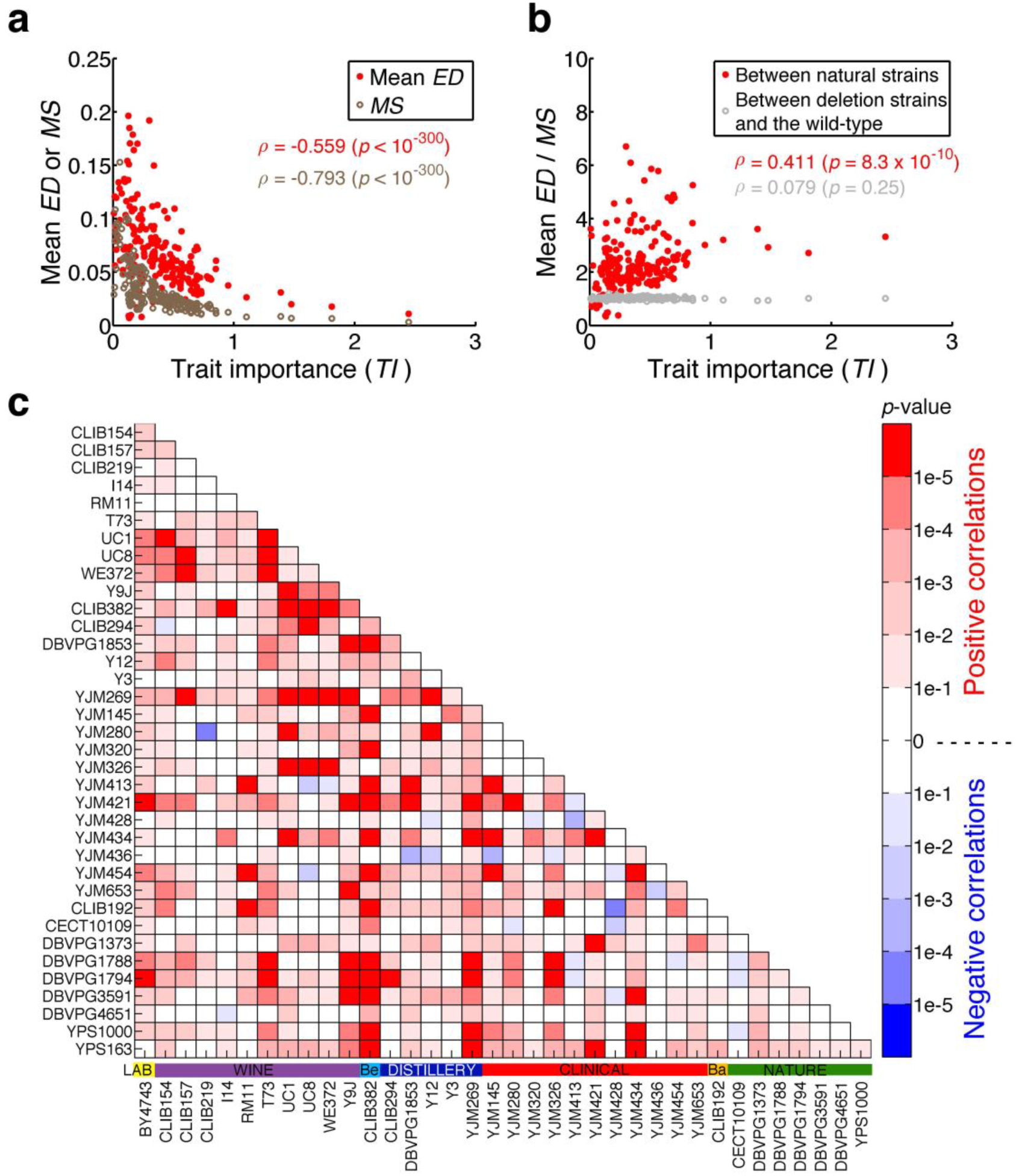
Prevalent adaptive evolution of morphological traits in the yeast *Saccharomyces cerevisiae*. **a**, Mean evolution distance (*ED*) of 666 pairs of natural strains for a trait and the mutational size (*MS*) of the trait both decrease with trait importance (*TI*). Each dot represents a morphological trait. ρ, Spearman’s rank correlation coefficient. **b,** Mean *ED* among 666 natural strain pairs for a trait relative to its *MS* increases significantly with *TI*, while the mean *ED* between 666 gene deletion strains and the wild-type relative to *MS* does not increase significantly with *TI*. **c,** Nominal *p*-values for the Spearman’s correlation between *ED*/*MS* and *TI* for all 666 pairs of natural strains. The horizontal colored bars above the strain names show the ecological environments of the strains. Be, beer; Ba, bakery.

To exclude the possibility that the above result is an artifact of our statistical analysis, we compared the wild-type BY strain with 666 randomly picked gene deletion strains of the BY background, under the premise that phenotypic differences between deletion strains and the wild-type should not be adaptive. As expected, there is no significant correlation between mean *ED*/*MS* and *TI* (Fig. 2b), where mean *ED* for a trait is the average phenotypic difference between these deletion strains and the wild-type for the trait.

Because the above *MS* and *TI* were estimated in haploid yeasts, while the 37 natural strains studied are diploid, we also estimated *MS* and *TI* using recently published morphological data of 130 diploid gene deletion strains^19^. We confirmed that the significant positive correlation between *ED*/*MS* and *TI* holds (**Extended Data Fig. 1**).

To examine whether the detected adaptive signal among the 37 natural strains is attributable to a small number of strains or is a general phenomenon of the species, we estimated the rank correlation (*ρ*) between *ED*/*MS* and *TI* for each of the 666 strain pairs, using the *ED* value of the strain pair. We found ρ to be positive for the vast majority of the strain pairs (Fig. 2c), suggesting pervasive adaptive morphological evolution in *S. cerevisiae*.

Some of the 210 morphological traits are genetically highly correlated^18,20^. To exclude the possibility that the adaptive signal is an artifact of the use of correlated traits, we estimated 210 principal component traits from the original traits (see Methods). Analysis of the principal component traits, which are independent from one another, shows even stronger signals of adaptive evolution (**Extended Data Fig. 2**).

If the prevalent positive ρ values among the 37 natural strains (Fig. 2c) truly arise from positive selection, we should expect that (i) the overall rate of morphological evolution is greater for strain pairs with higher ρ values and (ii) this rate disparity is primarily reflected in relatively important traits rather than relatively unimportant ones. We defined the overall rate of morphological evolution between two strains by their morphological dissimilarity across all 210 traits divided by their fractional genomic sequence difference. Consistent with our expectation, the rate of morphological evolution increases with ρ(*P* = 0.007, one-tail partial Mantel test with phylogenetic permutation to correct for both the nonindependence among strain pairs and phylogenetic relationships in the data; see Methods). Additionally, when we separated the 210 traits into two equal-sized bins based on *TI*, the correlation between the rate of morphological evolution and ρ remained significantly positive for the 105 traits with relatively high *TI* (*P* < 0.001) but not for the 105 traits with relatively low *TI* (*P* = 0.805). These observations support that, the greater the ρ value relative to 0, the stronger the positive selection on the morphological traits, especially relatively important ones.

What factors determine the ρ value of a strain pair? Because the 210 morphological traits were chosen purely based on experimental feasibility, we do not expect the detected adaptive signals to correlate with any obvious genetic or ecological factor. Indeed, using the partial Mantel test, we found no significant correlation between the ρ value of two strains and the strains’ difference in genome sequence, ecological environment, population membership, or geographic location (**Extended Data Table 1**). These findings are consistent with a previous analysis showing that the morphological similarities among the 37 natural strains cannot be explained by the strains’ similarities in population history or ecological environment^17^. Hence, the selective agents behind the detected morphological adaptations are unclear.

To test the neutral hypothesis of morphological evolution beyond the species level, we collected comparable morphological data from two strains of *S. paradoxus*, the sister species of *S. cerevisiae*, and one strain of their outgroup species *S. mikatae* (see Methods). We first calculated the mean *ED*/*MS* between the 37 *S. cerevisiae* strains and *S. paradoxus* strain N17 for each trait (**Supplementary Data 3**). Across the 210 traits, we observed a significantly positive correlation between mean *ED*/*MS* and *TI* (Fig. 3a). A similar pattern was observed between *S. cerevisiae* and *S. paradoxus* strain IFO1804 (Fig. 3b). Between *S. cerveisiae* and the more distantly related species of *S. mikatae* (**Supplementary Data 3**), however, the positive correlation is no longer significant (Fig. 3c), probably because of the lack of prevalent positive selection or a reduced statistical power as a result of using *MS* and *TI* values estimated from *S. cerevisiae* in tests involving distantly related species.

**Figure 3.**
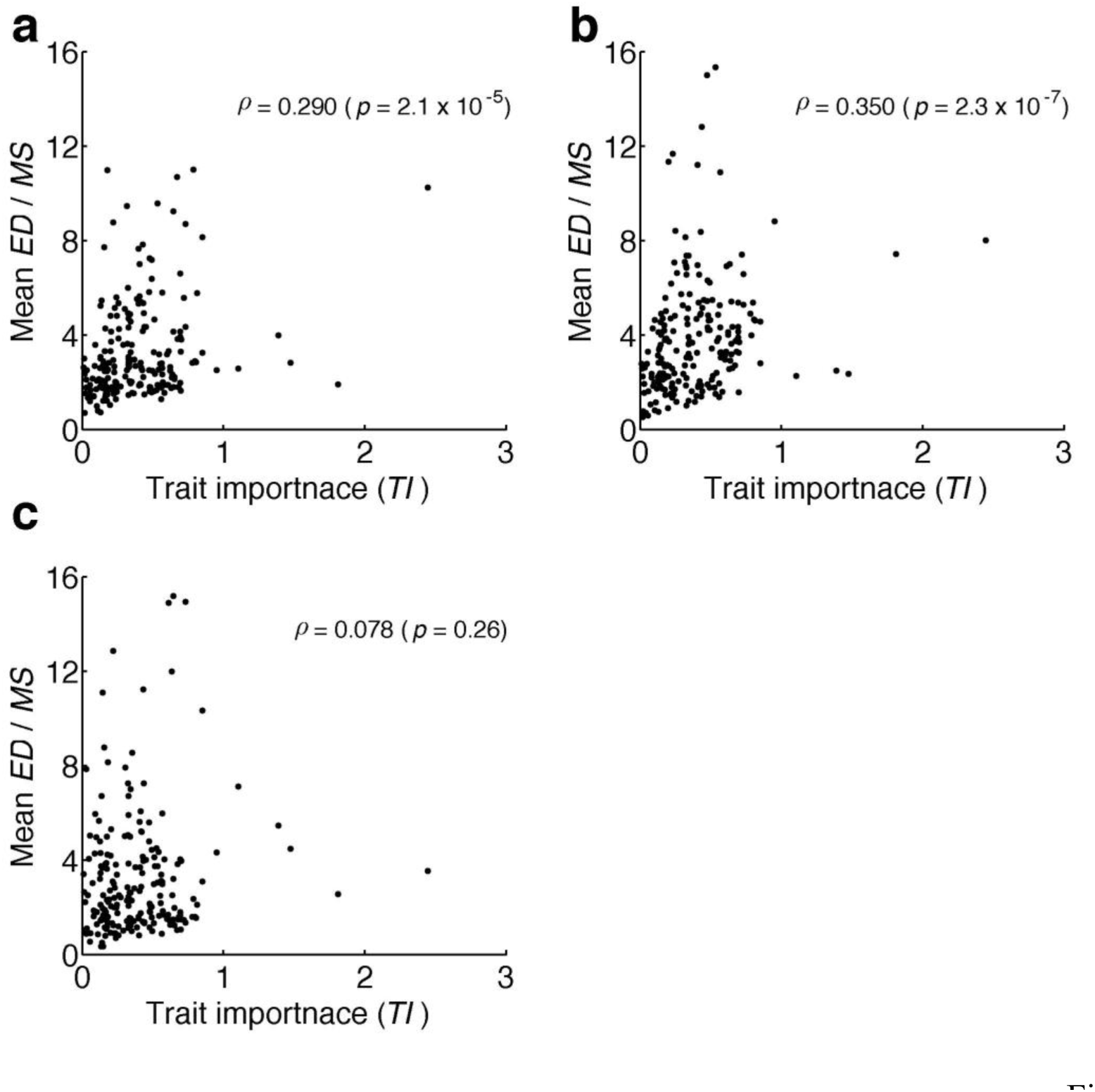
Adaptive morphological evolution between *Saccharomyces* species. **a**, The mean *ED*/*MS* between 37 natural *S. cerevisiae* strains and *S. paradoxus* strain N17 for a trait increases significantly with trait importance (*TI*).**b,** The mean *ED*/*MS* between 37 natural *S. cerevisiae* strains and the *S. paradoxus* strain IFO1804 increases significantly with *TI.* **c,** The mean *ED*/*MS* between 37 natural *S. cerevisiae* strains and *S. mikatae* increases (but not significantly) with *TI*. Each dot represents a morphological trait.

To investigate the generality of the above findings of adaptive phenotypic evolution, we turned to another class of traits that can be chosen regardless of their potential roles in adaptation: 3466 gene expression traits, each being the mRNA expression level of a yeast gene in a rich medium. Using microarray gene expression data, we quantified *ED* between two *S. cerevisiae* strains for each trait (**Supplementary Data 4**). We used two approaches to estimate *MS* on each trait. First, we estimated it from the microarray expression data of 1486 gene deletion strains, as in the case of morphological traits^18^. Second, we estimated it by the standard deviation (*SD*_m_) in relative expression level among four mutation accumulation lines^21^, arguably a more natural measure of mutational size. *TI* of the expression level of a gene was measured by the fitness reduction caused by the deletion of the gene^18^ (see Methods). Similar to the morphological data, gene expression data showed a negative correlation between *ED* and *TI* and a negative correlation between *MS* (or *SD*_m_) and *TI* (Table 1). However, contrary to the morphological traits, expression traits exhibited a significantly negative correlation between *ED*/*MS* (or *ED*/*SD*_m_) and *TI* (**Table 1**). While adaptive evolution of the expression levels of some yeast genes have been suggested^22-24^, overall we found relatively important genes to evolve more slowly in expression level than relatively unimportant ones, consistent with the neutral hypothesis. The similar results obtained from the use of *MS* and *SD*_m_ support that *MS* is a suitable proxy for mutational size. Here we are restricted to intra-specific analysis, because the use of different probes in the microarrays of different species prohibits a reliable comparison of the rate of expression evolution among genes. Use of other types of expression data such as mRNA sequencing data is currently infeasible, because no comparable data are available for estimating *MS*.

**Table 1.** Testing the neutral hypothesis of yeast gene expression evolution Variables correlated.

| **Variables correlated** | **Spearman's $\rho$** | **P-value** |
| --- | --- | --- |
| *ED, TI* | -0.147 | 3.7e-18 |
| *MS, TI* | -0.142 | 3.9e-17 |
| *ED/MS, TI* | -0.087 | 3.2e-07 |
| *SD<sub>m</sub>, TI* | -0.153 | 1.7e-19 |
| *ED/SD<sub>m</sub>, TI* | -0.060 | 4.5e-04 |

*ED*, evolutionary distance in gene expression level between two *S. cerevisiae* strains.
*TI*, gene expression trait importance.
*MS*, mutational size measured by gene deletion.
*SD<sub>m</sub>*, standard deviation in relative gene expression level among mutation accumulation lines.

In summary, our analysis of 210 yeast morphological traits with no *a priori* bias toward adaptive evolution reveals strong signals of adaptive evolution. To the best of our knowledge, this is the first demonstration of widespread adaptive evolution of a large set of random phenotypic traits in any organism. If these 210 traits are representative of yeast morphological traits, we must conclude that morphological evolution in yeast is frequently adaptive. Whether the same conclusion could be drawn in other species is unknown, but the methodology developed here is generally applicable, especially to genetic model organisms, in which mutational size estimation and trait importance estimation are relatively straightforward. Although it is essential to use a random set of traits to evaluate the prevalence of adaptive phenotypic evolution, using such traits means that the selective agent would be difficult to discern if adaptation is detected, as in the present case. Future work should attempt to identify the selective agents acting on these morphological traits, which are necessary for a complete understanding of phenotypic adaptation. It should be emphasized that, although the bar for rejecting neutrality is likely lower in our test than in most previous tests^3,4,11-13^, it remains quite high. This is because important traits that could be subject to strong positive selection are also expected to be under strong purifying selection such that an adaptive signal becomes detectable only when the difference in the strength of positive selection among traits surpasses that in the strength of purifying selection. The high bar renders claims of adaptation conservative. But, it also means that failure to reject neutrality, as in the case of yeast gene expression evolution, neither proves neutrality nor refutes adaptation. Nonetheless, based on additional experiments and tests, we recently found unambiguous evidence that the vast majority of gene expression level variations within and between yeast species are not adaptive but neutral^25^. Together, these analyses of yeast morphological and gene expression data raise the intriguing possibility that some classes of phenotypic traits evolve generally adaptively while others neutrally.

## Acknowledgements

We thank Seiko Morinaga for technical assistance and Xiaoshu Chen, Brian Metzger, and Jian-Rong Yang for valuable comments. This work was supported by U.S. National Institutes of Health research grant R01GM103232 to J.Z. and the Grants-in-Aid for Scientific Research grant 24370002 from the Ministry of Education, Culture, Sports, Science and Technology in Japan to Y.O.

**Extended Data Figure 1. Mean *ED* among 666 natural strain pairs of *S. cerevisiae* for a trait relative to its *MS* increases significantly with *TI*, when *MS* and *TI* are both estimated using diploid deletion strains.**

**Extended Data Figure 2. Prevalent adaptive evolution of morphological principal component traits in the yeast *S. cerevisiae*. a,** Mean evolution distance (*ED*) of 666 pairs of natural strains for a trait and the mutational size (*MS*) of the trait both decrease with trait importance (*TI*). Each dot represents a morphological principal component trait. ρ, Spearman’s rank correlation coefficient. **b,** Mean *ED* among 666 natural strain pairs relative to *MS* increases significantly with *TI*, while the mean *ED* between 666 gene deletion strains and the wild-type relative to *MS* does not increase significantly with *TI*. **c,** Nominal *p*-values for the Spearman’s correlation between *ED*/*MS* and *TI* for all 666 pairs of natural strains.

## METHODS

### A comparison between existing tests of the neutral hypothesis of phenotypic evolution and the newly proposed test

A number of neutrality tests of phenotypic evolution exist in the literature. They all test the neutrality of one trait and can be divided into two categories based on the rationale of the test. The first category compares the observed phenotypic distance in a trait between populations or species with the neutral expectation. Positive (or purifying) selection is inferred when the observed distance is significantly larger (or smaller) than the neutral expectation. The neutral expectation has been derived by considering the stochastic process of phenotypic evolution^3^ or genetic models of quantitative traits^4,13^. Various test statistics have been proposed, including for example Lande’s *N*_*e*_ ^3^, Lande’s *F* ^14^, Chakraborty and Nei’s *B*_*t*_/*V*_t_ ^13^, Turelli et al.’s MDE test^12^, and Lynch’s Δ ^15^ These tests usually require the information of effective population size and divergence time. They also require the information on the rate of mutational input such as narrow-sense heritability and mutational variance. If these parameters are unavailable, one may use an alternative approach by comparing *Q*_*ST*_ of a trait with *F*_*ST*_ of neutral loci, which are expected to be the same if the trait evolves neutrally^11^. Positive selection is undetectable by any of the above tests unless the intensity of positive selection exceeds that of purifying selection. Because purifying selection is expected to be pervasive even for traits subject to positive selection, all tests in this category have a low power in detecting positive selection.

The second category is the QTL sign test, which relies on the information from QTL mapping of a trait. Typically, positive selection is inferred when the number of positive-effect QTLs differs significantly from that of negative-effect QTLs^26^. Although this category of test is model-free, it cannot distinguish between positive selection and relaxation from purifying ^15^. These tests usually require the information of effective population size and selection without other information^23^.

Our test of the neutral hypothesis of phenotypic evolution compares the evolutionary rates of many traits with different levels of importance to fitness. Under the neutral hypothesis allowing purifying selection, relatively important traits evolve more slowly than relatively unimportant traits. If relatively important traits are found to evolve more rapidly than relatively unimportant ones, neutrality is rejected in favor of positive selection. Our test is expected to be more powerful than the above first category of tests, because detecting positive selection no longer requires the intensity of positive selection to exceed that of purifying selection. Rather, the criterion is that the difference in intensity between positive and negative selection is more positive for more important traits. Our test does not suffer from the problem in the above second category of tests, because it is extremely improbable for the relative importance of a large number of traits to be reversed in a second population, compared with that in the original population where trait importance is measured.

The yeast morphological data from natural strains and gene deletion strains, gene expression data, and the fitness data of gene deletion strains used for estimating trait importance were all collected under a laboratory rich medium. While this medium does not equal the many natural environments of various yeast strains, this discrepancy does not affect our neutrality test, because all required by our test is that the same condition is used in phenotyping a trait and in estimating its trait importance. The main problem with mismatches between the experimental condition and the natural environments is that the selective agent becomes more difficult to discern.

### Yeast morphological data

The morphological traits analyzed here were previously defined^16^. Briefly, yeast cultures were grown to 1×10^7^ cells/ml in YPD or synthetic complete media. Cells were fixed with 3.7% formaldehyde and stained with fluorescein isothiocyanate-Con A, rhodaminephalloidin, and 4 diamidino-2-phenylindole, which simultaneously mark cell wall, actin cytoskeleton, and nuclear DNA, respectively. After digital images were acquired, cell images were collected and processed by *CalMorph*. In total, 501 morphological traits were measured. Among these morphological traits, we focused on 210 traits, which are defined for individual cells rather than cell populations, have positive trait importance values^18^, and were measured in all the strains analyzed here. The morphological data of *S. paradoxus* strain N17, *S. paradoxus* strain IFO 1804, and *S. mikatae* strain IFO1815 were generated with 15, 10, and five biological replicates, respectively. On average, ∼60 cells were measured for each trait in each replicate. The morphological data from 37 natural strains of *S. cerevisiae* were previously collected, with five biological replicates per strain^17^. The morphological data of 4718 haploid and 130 diploid single gene deletion strains of *S. cerevisiae* were previously published^16,19^.

### Evolutionary distance between two strains for a morphological trait

We estimated the mean phenotypic value for a trait in a strain by first calculating the mean trait value among all cells in a replicate population and then averaging this number across all replicate populations. Let *x*_i_ and *x*_j_ be the mean phenotypic values of a trait in strains *i* and *j,* respectively. We estimated the raw evolutionary distance for the trait between strains *i* and *j* by *ED*_ij_ = |*x*_i_ - *x*_j_| / [(*x*_i_ + *x*_j_)/2]. Different traits have different levels of environmental variation in phenotyping, different levels of random measurement error, and different levels of among-individual stochastic phenotypic variation. To allow a comparison among traits, we corrected the above estimated raw *ED* values for these factors ^18^. Let us assume that strain *i* has *m* replicate populations, with population sizes of *a*_1_, *a*_2_,…, *a*_m_, respectively, and strain *j* has *n* replicate populations, with population sizes of *b*_1_, *b*_2_,…, *b*_n_, respectively. We generated 100 sets of bootstrap samples for both strain *i* (i.e., each having *m* populations with sizes of *a*_1_, *a*_2_,… *a*_m_) and strain *j* (i.e., each having *n* populations with sizes of *b*_1_, *b*_2_,…, *b*_n_) using the data from the *n* populations of strain *j*. Note that the hierarchical structure of the data is retained in bootstrapping. We then estimated the average *ED*_ij_ across these 100 sets of bootstrapped samples, and denoted it by pseudo *ED*_j_→_i_. We similarly estimated pseudo *ED*_i_→_j_ using the data from the *m* populations of strain *i*. We then averaged *ED*_j→i_ and *ED*_i→j_ and subtracted this value from raw *ED*_ij_ to obtain the corrected *ED*_ij_. If the corrected *ED*_ij_ is negative, we set it at 0. All *ED* values presented in the main text and figures are corrected *ED*_ij_.

### Mutational size

The mutational size for a trait was calculated by the average of the previously published net effect sizes of 4718 haploid single gene deletions on the trait^18^. Briefly, we first calculated the raw effect size (*ES*_ij_) of deleting gene *i* on trait *j* as (*x*_ij_ - *w*_j_)/*w*_j_, where *x*_ij_ is the mean phenotypic value of trait *j* in the deletion strain *i*, and *w*_j_ is the corresponding value in the wild-type (averaged across 123 replicate populations). Then we generated 1000 pseudo phenotypic datasets to estimate the net |*ES*| of gene deletion on a trait. To generate a pseudo dataset, we randomly chose one of the 123 wild-type replicate populations and picked (with replacement) from this population the same number of cells as in the actual gene-deletion data. We then calculated mean pseudo |*ES*| across all pseudo datasets, and net |*ES*| equals raw |*ES*| minus mean pseudo |*ES*| if raw |*ES*| > mean pseudo |*ES*| or zero if raw |*ES*| < mean pseudo |*ES*|.

We also estimated mutational size by the same method but using the recently published morphological data of 130 diploid single gene deletion strains^19^ along with the five replicate populations of the corresponding wild-type (BY4743)^17^. Due to the low number of wild-type replicates, 100 pseudo datasets were generated.

### Morphological trait importance

The trait importance (*TI*) of each of the 210 morphological traits was previously estimated from the negative slope of the linear regression between the corrected phenotypic effect of a gene deletion on the trait and the fitness of the gene deletion strain across 2779 haploid single gene deletion strains^18^. The deletion strains with fitness larger than 1 were not used. Briefly, for each trait *j*, we performed a linear regression *F*_i_ = *a*_j_ - *b*_j_ (net |*ES*_ij_|), where net |*ES*_ij_| is the absolute value of the net effect size of deleting gene *i* on trait *j*, and *F*_i_ is the fitness of the strain lacking gene *i* relative to the wild-type in YPD. In this regression, the estimated slope *b*_j_ > 0 is 100 times the reduction in fitness caused by 1% change in the phenotypic value of trait *j*, while the estimated intercept *a*_j_ is the expected fitness when net |*ES*_ij_| = 0. Thus, *b*_j_ is a measure of the relative importance of trait *j* to fitness, or trait importance (*TI*).

Because *TI* is estimated by the correlation between phenotypic changes and fitness changes, it may not accurately reflect the causal relationship between the variation of a trait and fitness. The fact that our use of the inaccurate *TI* estimates still yields significant evidence for adaptive morphological evolution suggests that the true signal is even stronger. In other words, our results are likely to be conservative.

For diploid single gene deletion strains^19^, we applied the same method to calculate *TI*, but in this case only 99 strains with fitness ≤ 1 could be used.

### Principal component analysis

To examine if the non-independence among the 210 morphological traits affects our results, we followed previous studies^18,20^ to perform a principal component analysis to transform the net |*ES*| matrix **M** (4718 genes × 210 traits) described previously^18^. Note that Net |*ES*| is the corrected absolute effect size of a gene deletion on a trait. After this function returned a coefficient matrix **C** (210 × 210), we calculated ***v***’ = ***v* C** (1 × 210 principal traits) for each corrected *ED* vector *****v***** (1 × 210 traits) between two strains. The absolute values of ***v***’ provided the corrected *ED* for each of the 210 orthogonal principal component traits. We also used the transformed net |*ES*/ matrix, **M’=MC** (4718 genes × 210 principal traits), to estimate trait importance for the 210 principal component traits following the method previously used^18^.

### Mantel’s test

We used Mantel’s test^27,28^ to evaluate the significance of the correlation between a biological distance (e.g., morphological dissimilarity between two strains) and the correlation (ρ) between *ED*/*MS* and *TI* across 666 pairs of the 37 natural strains of *S. cerevisiae*. In general, Mantel’s test evaluates whether the correlation between two distance matrices is significant by comparing the test statistic (*z*) calculated by the observed matrices and the null distribution of *z* calculated by randomly shuffling one of the matrices. The test statistic could be an element-by-element product or a Pearson correlation coefficient between all elements. During each shuffling, two columns are randomly picked and swapped, and the corresponding rows are also swapped. In this study, we used the correlation coefficient between all elements of two matrices as the test statistic and generated the null distribution of the test statistic by randomly shuffling one of the matrices 1000 times.

When the dataset has underlying phylogenetic relationships such as the present dataset in which all strains are connected by a phylogeny, the significance level may be inflated because of the shared evolutionary history among lineages^29^. To control the phylogenetic non-independence, we performed a partial Mantel test, in which the phylogenetic distance matrix is controlled^30^. Among four different ways to implement the partial Mantel test, we used the so-called method 2, in which the residual matrix instead of the original matrix is shuffled, as suggested^31^. In addition, we used phylogenetic permutation (PP) in partial Mantel test to obtain non-inflated type-I errors^32^. Specifically, applying PP means that the chance to pick a strain pair to swap is proportional to their phylogenetic distance^33^. The phylogenetic tree of the 37 natural strains of *S. cerevisiae* was reconstructed using the single nucleotide polymorphism (SNP) data from Maclean et al.^34^ by the neighboring-joining method^35^ with *p*-distance^36^ implemented in MEGA 5.2.2 ^37^, and the matrix of pairwise phylogenetic distances was obtained by APE in R ^38^.

We used the Spearman’s correlation coefficient (ρ) between *ED/ES* and *TI* of a pair of strains as a measure of adaptation. By partial Mantel’s test, we examined whether ρ is correlated with the following five parameters for the strain pair: morphological evolutionary rate, dissimilarity in genome sequence, dissimilarity in ecological environment, dissimilarity in population membership, and dissimilarity in geographic location. The morphological evolutionary rate between two strains was estimated as follows. First, for each trait, we ranked the 37 strains by their mean phenotypic values for the trait. Second, we calculated Spearman’s correlation between the 210 ranks of one strain with those of the other strain. The rate of morphological evolution was calculated by the negative of this Spearman’s correlation divided by genomic sequence dissimilarity (see below). Genomic sequence dissimilarity was calculated by the SNP density between two strains^34^. Ecological environment dissimilarity was set as 0 if two strains were sampled from the same ecological environment and 1 if sampled from different environments^34^. Population membership dissimilarity was set as 0 if two strains belong to the same population based on the fastSTRUCTURE analysis of the SNP data of 190 yeast strains^34^; otherwise, it was set as 1. Strains designated as “mosaics”^34^ were excluded from this analysis because mosaics do not represent a population. Geographic location dissimilarity was set to 0 if two strains were sampled from the same continent; otherwise, it was set to 1. Note that CLIB382 was removed from this analysis because of the lack of SNP data in Maclean et al.^34^. Furthermore, any strain lacking information for any parameter was removed from the analysis for that particular parameter.

### Gene expression data and analysis

The rich medium microarray gene expression ratio (*r*) between two *S. cerevisiae* strains, RM and BY, were previously measured for thousands of genes^39^. We defined the intra-specific evolutionary distance (*ED*) of the expression level for a gene by |*x*_RM_-*x*_BY_|/*x*_BY_, which equals |*r*-1|, because *r* = *x*_RM_/*x*_BY_, where _RM_ and _BY_ are expression levels of the gene in *x*_RM_ and *x*_BY_, respectively.

We used two different methods to estimate mutational size for each expression trait. The first method was to use the mean effect size of gene deletion on the expression trait across 1486 deletion lines, following a previously published method^18^ but using a recently published large dataset^40^. The second method was to use the square root of variance in the expression level of an evolved line relative to that of the ancestral line (*SD*_m_) among four mutation accumulation lines^21^. The trait importance (*TI*) of a gene expression trait was defined by the fitness decrease caused by deleting the gene^22^, and only those genes that cause a fitness reduction of zero or more when deleted were considered. After removing genes that miss any kind of data above, we obtained our final dataset with 3466 expression traits.

